# *Kif11*-haploinsufficient oocytes reveal spatially differential requirements for chromosome biorientation in the spindle

**DOI:** 10.1101/2025.01.02.631055

**Authors:** Tappei Mishina, Aurélien Courtois, Hiroshi Kiyonari, Tomoya S. Kitajima

**Affiliations:** Laboratory for Chromosome Segregation, RIKEN Center for Biosystems Dynamics Research (BDR), Kobe, Japan; Faculty of Agriculture, Kyushu University, Fukuoka, Japan; Laboratory for Animal Resources and Genetic Engineering, RIKEN Center for Biosystems Dynamics Research (BDR), Kobe, Japan; Sex Chromosome Biology Laboratory, The Francis Crick Institute, London, UK

## Abstract

Bipolar spindle assembly and chromosome biorientation are prerequisites for chromosome segregation during cell division. The kinesin motor KIF11 drives spindle bipolarization by sliding antiparallel microtubules bidirectionally, elongating a spherical spindle into a bipolar-shaped structure in oocytes. This process stretches chromosomes, establishing chromosome biorientation at the spindle equator. The quantitative requirement for KIF11 in spindle bipolarization and chromosome biorientation remains unclear. Here, using a genetic strategy to modulate KIF11 expression levels, we show that *Kif11* haploinsufficiency impairs spindle elongation, leading to the formation of a partially bipolarized spindle during meiosis I in mouse oocytes. While the partially bipolarized spindle allows chromosome stretching in the inner region of its equator, it fails to do so in the outer region. These findings demonstrate the necessity of biallelic functional *Kif11* for bipolar spindle assembly in oocytes and reveal a spatially differential requirement for chromosome biorientation within the spindle.

## Introduction

Accurate chromosome segregation during female meiosis is indispensable for the faithful transmission of genetic information to the next generation. Errors in this process result in the production of aneuploid eggs, fertilization of which causes pre-implantation loss, miscarriage, or congenital disorders such as Down syndrome (Herbert *et al*, 2015; Charalambous *et al*, 2023).

Chromosome segregation is driven by the spindle, a microtubule-based dynamic machine. Entry to the M-phase of meiosis I promotes microtubule polymerization depending on chromosome-derived diffusible signals (Schuh & Ellenberg, 2007; Holubcová *et al*, 2015; Dumont *et al*, 2007; Drutovic *et al*, 2020). During prometaphase, in mammalian oocytes, these microtubules initially assemble into an apolar spherical spindle (Schuh & Ellenberg, 2007). Subsequently, KIF11, a plus-end-directed kinesin, crosslinks antiparallel microtubules and slides them bidirectionally (Kapitein *et al*, 2005). This activity drives progressive elongation of the spindle, transforming it into a bipolar-shaped structure during the prometaphase-to-metaphase transition (Schuh & Ellenberg, 2007). Spindle elongation allows microtubules to stretch chromosomes toward the opposite poles (Schuh & Ellenberg, 2007; Kitajima *et al*, 2011). Stretched chromosomes align at the spindle equator, establishing chromosome biorientation and forming a spatial arrangement called the metaphase plate. Although both spindle bipolarization and chromosome biorientation are KIF11-dependent processes required for proper chromosome segregation, the spatiotemporal coordination between these processes remains poorly understood.

Chromosome dynamics within the spindle are spatially inhomogeneous. In both somatic cells and oocytes, chromosome oscillations are more pronounced in the inner region of the spindle equator (Civelekoglu-Scholey *et al*, 2013; Takenouchi *et al*, 2024). In addition, in oocytes, chromosomes located in the inner region of the spindle equator stretch earlier than those in the outer region (Takenouchi *et al*, 2024). Furthermore, stretched chromosomes in the inner region exhibit increased centromere-to-kinetochore distances (Takenouchi *et al*, 2024), indicative of stronger microtubule-mediated pulling forces. These observations may imply that the spindle produces stronger KIF11-dependent bipolar pulling forces in the inner region of its equator. However, it remains unknown whether the amount of KIF11 required for chromosome stretching differs between the inner and outer regions of the spindle equator.

Elucidating the quantitative requirements for KIF11 is critical for advancing molecular understanding underlying female infertility. A recent study has identified heterozygous mutations in the *Kif11* gene in patients experiencing repeated failures of *in vitro* fertilization or intracytoplasmic sperm injection at fertility clinics (Wu *et al*, 2024). Overexpression of these mutant forms of KIF11 has been shown to exert dominant-negative effects, disrupting efficient chromosome alignment (Wu *et al*, 2024). However, whether a heterozygous loss-of-function mutation of *Kif11* impairs spindle bipolarization or chromosome biorientation remains unknown.

In this study, we establish a gene knockout strategy to investigate the quantitative requirements for KIF11 in mouse oocytes. We show that heterozygous knockout of the *Kif11* gene in oocytes compromises spindle elongation during the prometaphase-to-metaphase transition of meiosis I, followed by a significant delay or block in anaphase onset. *Kif11*-haploinsufficient oocytes exhibit a partially bipolarized spindle, with chromosomes stretching in the inner region of the metaphase plate but not in the outer region. These results demonstrate that the biallelic expression of functional KIF11 is required for oocyte meiosis and reveal a spatially differential nature of chromosome biorientation on the spindle equator.

## Results

### Gene knockout strategy to study the quantitative requirements for KIF11 in mouse oocytes

To investigate the quantitative requirements for KIF11 in mouse oocytes, we used a strategy with heterozygous constitutive deletion and meiotic-specific conditional deletion of the *Kif11* gene (Fig. 1A, B). To generate the conditional deletion, we crossed our newly established floxed *Kif11* allele (*Kif11^fl^,* Fig. S1A) mouse line with the meiosis-specific *Spo11-*Cre recombinase mouse line (Lyndaker *et al*, 2013). Offspring derived from mice carrying the conditional deletion allele were used to establish a constitutive deletion allele of *Kif11* (*Kif11^del^*). We estimated the expression levels of full-length Kif11 mRNA in isolated fully-grown oocytes using poly(A) RNA sequencing followed by counting of reads corresponding to the first exon of the *Kif11* gene (Fig. 1C). This analysis showed that heterozygous conditional deletion (*Spo11-Cre, Kif11^wt/fl^*) reduced Kif11 levels to 59.5%, whereas heterozygous constitutive deletion (*Kif11^fl/del^*without *Spo11-Cre*) reduced them to 22.1% (Fig. 1D). Heterozygous combination of the conditional and constitutive deletion alleles (*Spo11-Cre, Kif11^fl/del^*) resulted in a reduction of Kif11 levels to only 3.0% of the wild-type expression levels (Fig. 1D).

**Figure 1:**
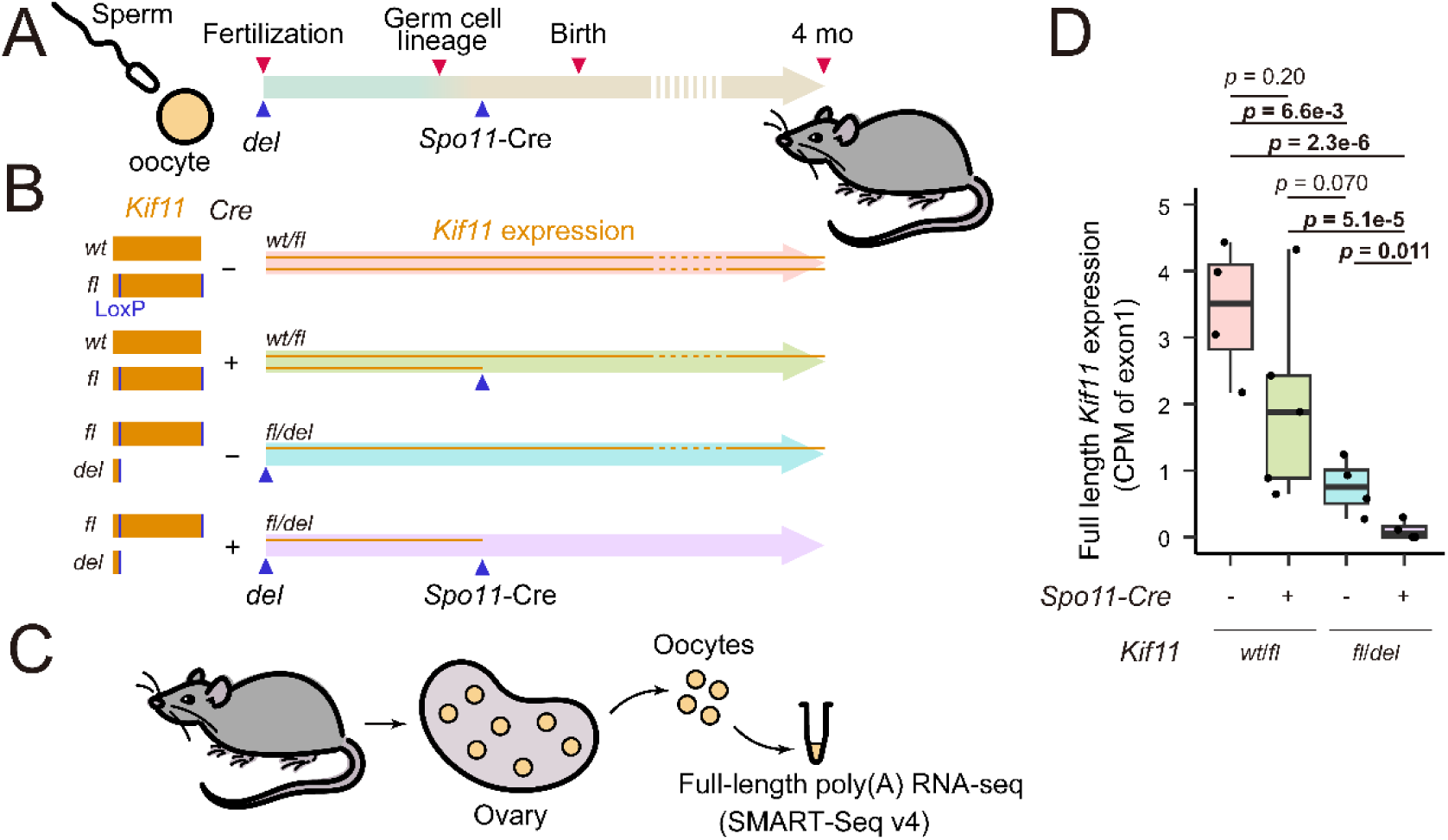
Gene knockout strategy to study the quantitative requirements for KIF11 in mouse oocytes. (A,B) Schematic of the generation of *Kif11* floxed mice. (A) *Spo11*-Cre allows activation of Cre in early prophase. (B) duration of *Kif11* expression under different genotypes of the combination of *Kif11* (wild-type, floxed, and deleted allele) and *Spo11*-Cre. The orange lines within arrows the indicate the expression of *Kif11*. (C) Design overview of the low input full-length poly(A) RNA sequencing using SMART-Seq v4 technology. (C) Box plot comparing the relative expression levels of full-length *Kif11* transcripts as measured by the coverage of the first exon from oocytes of 4-month-old mice oocytes with respect to their genotype (Holm’s correction for multiple comparisons; oocytes pooled from three or four independent experiments)

To verify the depletion of KIF11 protein, we used single-oocyte immunofluorescence assay, given the unavailability of antibodies allowing a reliable detection of KIF11 by Western blotting on the limited amount of oocyte extracts. Quantification of immunofluorescence signals at metaphase of meiosis I showed that KIF11 levels on the spindle tended to be reduced in heterozygous conditional deletion oocytes (*Spo11-Cre, Kif11^wt/fl^*) and in heterozygous constitutive deletion oocytes (*Kif11^fl/del^* without *Spo11-Cre*) (Fig. S2A and B), whereas the heterozygous combination of the conditional and constitutive deletion alleles (*Spo11-Cre, Kif11^fl/del^*) resulted in a nearly complete depletion of KIF11 signals (Fig. S2A and B).

We then assessed chromosome alignment on these images. Quantification of the spatial distribution of chromosomes (Fig. S2C) showed that whereas chromosomes were aligned on the metaphase plate in control oocytes, misaligned chromosomes were significantly increased in heterozygous constitutive deletion (*Kif11^fl/del^* without *Spo11-Cre*) oocytes (Fig. S2D). Chromosome alignment was never observed in oocytes of the heterozygous combination of conditional and constitutive deletion (*Spo11-Cre, Kif11^fl/del^*) (Fig. S2D). Thus, the genetically modified oocytes established in this study can serve as models to investigate dose-dependent requirements for KIF11 in oocyte meiosis.

### Spindle elongation requires a full dose of KIF11

To monitor spindle bipolarization, we performed live imaging of the microtubule marker EGFP-MAP4 and the chromosome marker H2B-mCherry throughout M-phase of meiosis I (Schuh & Ellenberg, 2007; Kitajima *et al*, 2011) (Fig. 2A, Movie EV1). Quantification of spindle shape in 3D showed that control oocytes initially formed an apolar spherical spindle, which gradually elongated into a bipolar-shaped structure (Fig. 2A; Movie EV1), as indicated by a progressive increase in its aspect ratio (Fig. 2B, C). In contrast, spindle elongation was severely perturbed in heterozygous conditional deletion oocytes (*Spo11-Cre, Kif11^wt/fl^*) and in heterozygous constitutive deletion oocytes (*Kif11^fl/del^* without *Spo11-Cre*) (Fig. 2A, C; Movie EV1). These oocytes subsequently exhibited a severe delay or block in anaphase entry (Fig. 2D). Oocytes with the heterozygous combination of conditional and constitutive deletions (*Spo11-Cre, Kif11^fl/del^*) exhibited almost no detectable spindle elongation or anaphase entry (Fig. 2A, C, D; Movie EV1). These results demonstrate that efficient spindle elongation and anaphase entry require the biallelic expression of functional KIF11.

**Figure 2:**
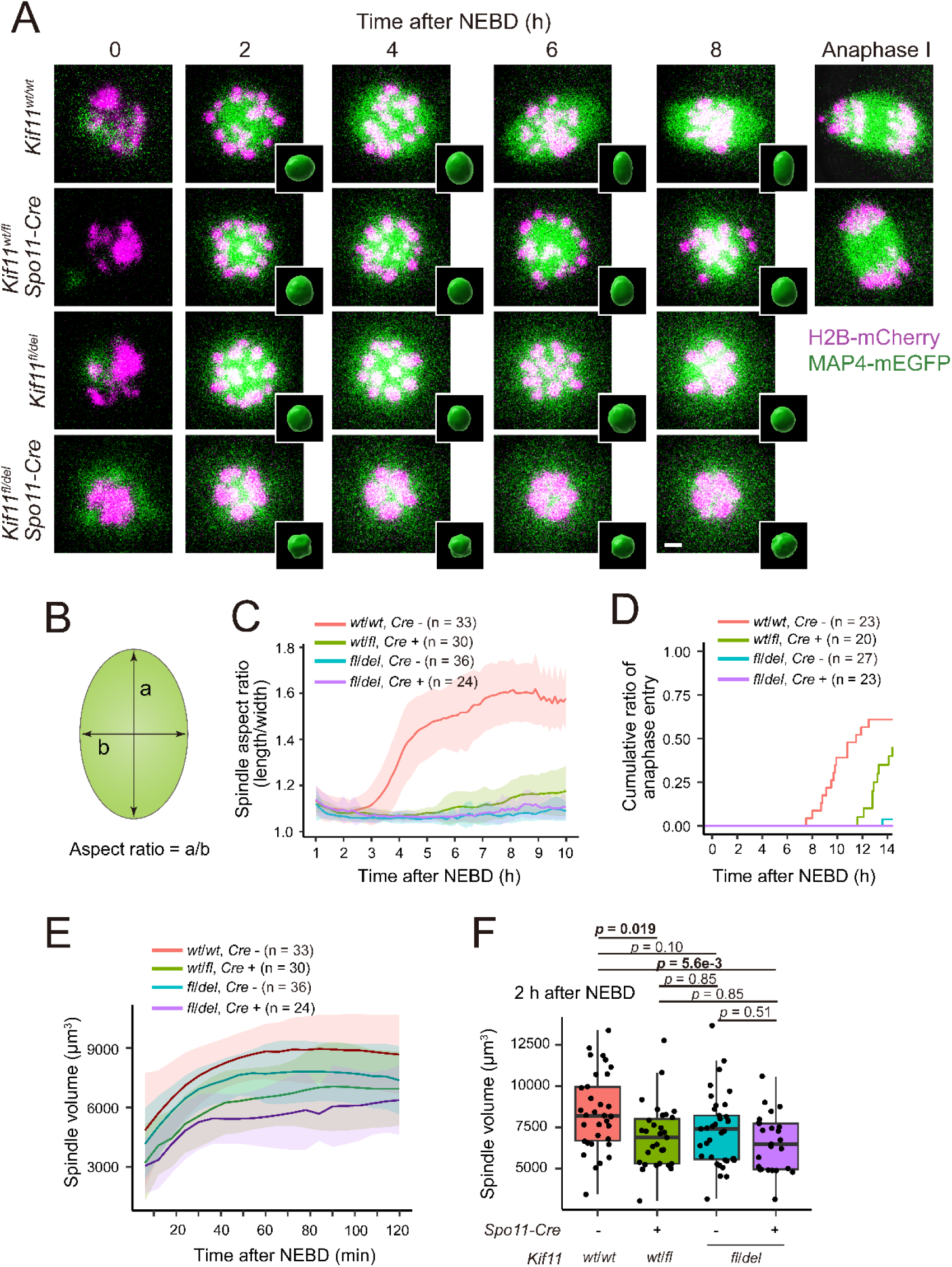
Spindle elongation requires a full dose of KIF11. (A) Live imaging of *Kif11^wt/wt^* (control), *Kif11^wt/fl^ Spo11-Cre* (single allele depletion from early prophase of meiosis), *Kif11^fl/del^* (complete single allele depletion), and *Kif11^fl/del^ Spo11-Cre* oocytes expressing EGFP-MAP4 (microtubules, green) and H2B-mCherry (chromosomes, magenta). Z-projection images are shown. 3D-reconstructed spindles are shown in insets. Scale bars 5 µm. (B) Diagram for measuring the aspect ratio (length/width) of the 3D-reconstructed spindle. (C) Quantification of spindle bipolarization shown as mean ± SD. Aspect ratio was measured. (D) Timing of anaphase entry for each genotype. (E) Spindle volume over time with mean ± SD. Spindle volume was determined from 3D-reconstructed images. (F) Box plot comparing spindle volume at prometaphase (2 hours after nuclear envelope breakdown, NEBD) between genotypes (Welch’s t-test comparing all possible pairs with Holm’s correction for multiple comparisons). Three (C, E, F) or two (D) independent experiments were performed.

We also noticed that KIF11-insufficient oocytes had a significantly smaller spindle volume, which was evident already before the time when control oocytes initiated spindle elongation (2 hours after nuclear envelope breakdown, NEBD) (Fig. 2A, E, F; Movie EV1). In contrast, treatment of oocytes with monastrol, an inhibitor of KIF11 motor activity (Kapoor *et al*, 2000), did not significantly reduce the volume of the microtubule mass (Fig. S3A–D). These results are consistent with the idea that KIF11 protein, but not its motor activity, promotes microtubule polymerization in the early stage of spindle formation.

### *Kif11*-haploinsufficient oocytes form a partially bipolarized spindle

Although *Kif11*-haploinsufficient spindles lack an elongated shape, it is still possible that they have bipolar microtubule arrays. To address this, we used mNeonGreen-CEP192, a marker for acentriolar microtubule-organizing centers (MTOCs), which are located at spindle poles where microtubule minus ends are enriched (Clift & Schuh, 2015). As expected, live imaging showed that CEP192-labeled MTOCs were progressively enriched at spindle poles in control oocytes (Fig. 3A, Movie EV2). We found that heterozygous conditional deletion oocytes (*Spo11-Cre, Kif11^wt/fl^*) also exhibited a bipolar distribution of MTOCs (Fig. 3A–C; Movie EV2), although the distance between the two poles appeared to be smaller than that of control oocytes (Fig. 3D). In heterozygous constitutive deletion oocytes (*Spo11-Cre, Kif11^fl/del^*), the bipolar distribution of MTOCs was severely defective, although a weak bipolar enrichment was occasionally observed (Fig. 3A–C; Movie EV2). In oocytes with the heterozygous combination of conditional and constitutive deletions (*Spo11-Cre, Kif11^fl/del^*), no bipolar MTOC distribution was observed (Fig. 3A–C; Movie EV2). These results suggest that the spindle in *Kif11*-haploinsufficient oocytes, while failing to elongate, manages to establish a partial bipolarity.

**Figure 3:**
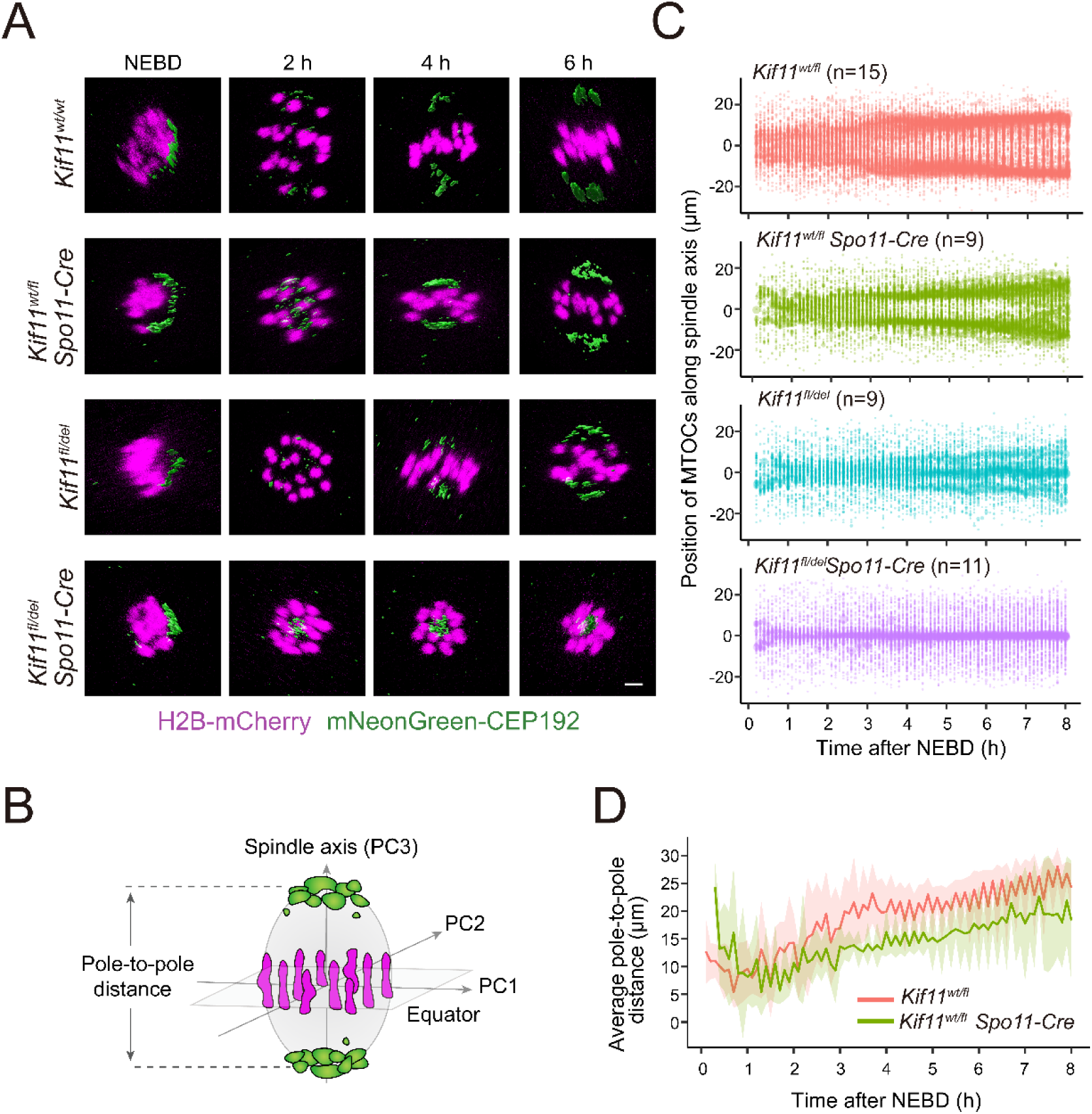
*Kif11*-haploinsufficient oocytes form a partially bipolarized spindle. (A) Live imaging oocytes expressing mNeonGreen-CEP192 (MTOCs, green; surface rendered) and H2B-mCherry (chromosomes, magenta) for each *Kif11* deletion genotype. 3D-reconstructed images with signal interpolation in z are shown. Scale bars 5 µm. (B) Diagram showing determination of the spindle axis and equator and the measurement of pole-to-pole distances based on chromosome and MTOC positions. (C) Bipolar MTOC sorting depends on KIF11 dosage. MTOC volume and its position along the spindle axis over time were measured from 3D-reconstructed images. The dots represent each MTOC with its size scaled by the relative MTOC volume. (D) Line plot comparing the two pole-to-pole distances between *Kif11^wt/fl^*and *Kif11^wt/fl^ Spo11-Cre* oocytes. Two independent experiments were performed.

### Concentric pattern of KIF11 dosage dependency for chromosome biorientation on the spindle equator

These findings led us to investigate how KIF11 insufficiency affects chromosome biorientation. In control oocytes, all chromosomes established a stretched state at the spindle equator by metaphase, as indicated by an increased distance between homologous kinetochores (inter-kinetochore distance) (Fig. 4A, B). No spatial bias in inter-kinetochore distance on the spindle equator was observed in control oocytes (Fig. 4C, D), consistent with our previous report (Takenouchi *et al*, 2024). Interestingly, we found that in heterozygous conditional deletion oocytes (*Spo11-Cre, Kif11^wt/fl^*), chromosomes in the inner region of the spindle equator were stretched with an increased inter-kinetochore distance, indicative of chromosome biorientation, while chromosomes in the outer region were not (Fig. 4A–D). In heterozygous constitutive deletion oocytes (*Spo11-Cre, Kif11^fl/del^*), most chromosomes were not stretched and were preferentially located in the outer region of the spindle equator (Fig. 4A–D). These results suggest that a higher dose of KIF11 is required for chromosome biorientation at the outer region of the spindle equator.

**Figure 4:**
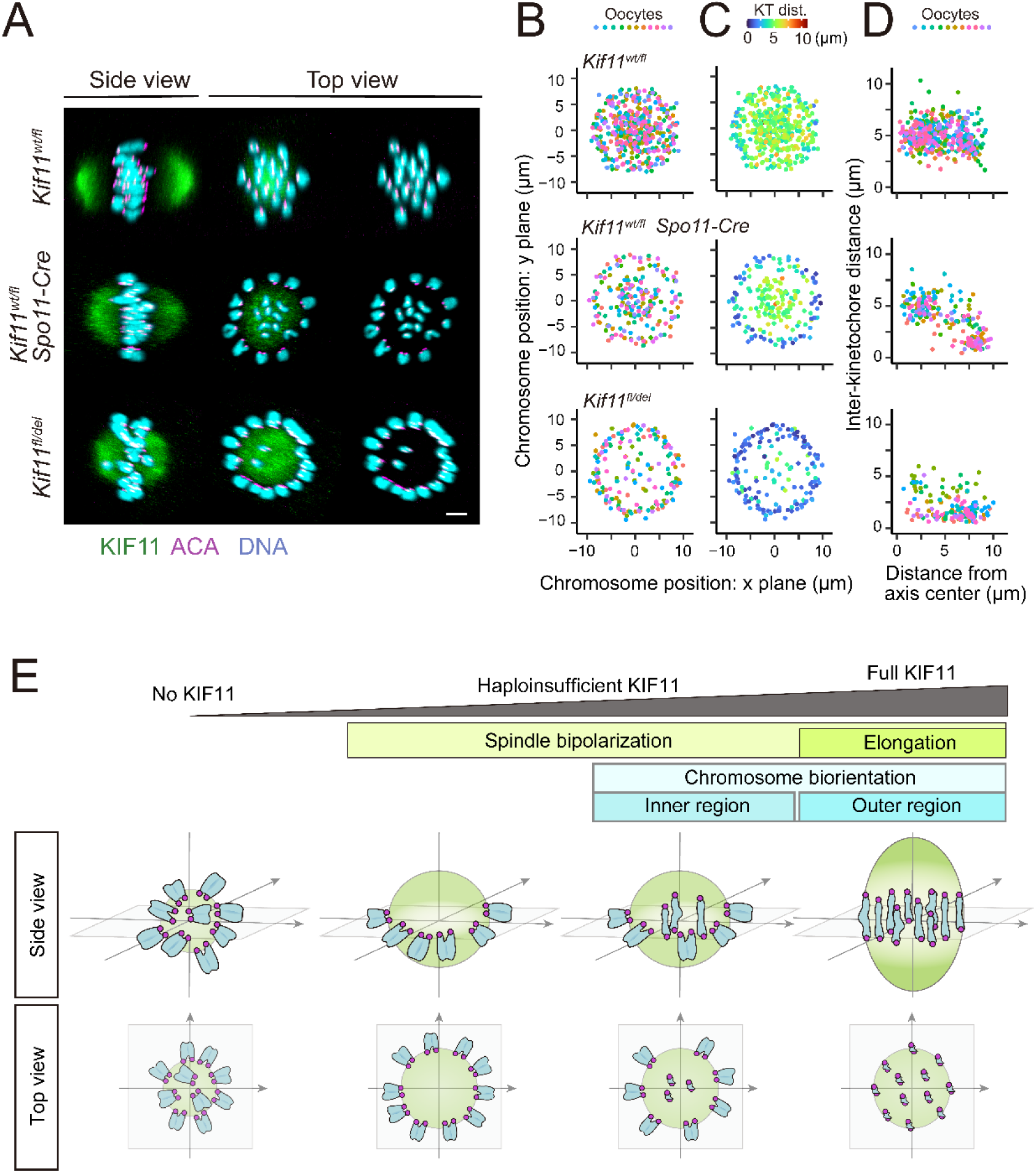
Concentric pattern of KIF11 dosage dependency for chromosome biorientation on the spindle equator. (A) Representative images of chromosome position and KIF11 localization for each genotype. Images are viewed in 3D with signal interpolation in z. Scale bar, 5 μm. (B) Distribution of chromosome positions on the spindle equator is shown. Each dot represents a chromosome. Colors indicate oocytes. (C) Chromosome positions are shown as in B, with color codes indicating inter-kinetochore distance. (D) 2D plot of inter-homologous kinetochore distance with respect to its position from the center of the spindle equator (right). Colors indicate oocytes. (E) Spatially differential requirement of KIF11 dose for chromosome biorientation within the spindle. In oocytes with haploinsufficient KIF11, spindle elongation is impaired, leading to the formation of a partially bipolarized spindle, where chromosomes biorient in the inner region of the spindle equator but fail to do so in the outer region. Two independent experiments were performed.

## Discussion

In this study, we established mouse oocyte models with insufficient expression levels of KIF11, demonstrating that *Kif11* haploinsufficiency causes severe spindle defects. Furthermore, our results reveal a spatially differential requirement for chromosome biorientation on the spindle equator: while the reduced levels of KIF11 are sufficient for chromosome stretching in the inner region, a full dose of KIF11 is required in the outer region (Fig. 4E).

We propose two possible explanations for the spatially differential requirement for chromosome biorientation. One possibility is that the polar ejection force is stronger in the outer region of the spindle equator. This force, which pushes chromosomes from the spindle poles toward the equator, can counteract KIF11-dependent bipolar microtubule pulling forces required for chromosome biorientation (Rieder *et al*, 1986; Chong *et al*, 2024). Stronger polar ejection forces in the outer region of the spindle equator may also explain less pronounced chromosome oscillation, as suggested in studies using somatic cells (Civelekoglu-Scholey *et al*, 2013).

Alternatively, factors that cooperate with KIF11 to promote chromosome biorientation may be more concentrated in the inner region of the spindle equator. Being occupied and surrounded by chromosomes, the inner region may experience higher concentrations of chromosome-derived signals, creating a microenvironment that enhances KIF11-mediated chromosome biorientation. For instance, chromosomes produce diffusible RanGTP signals that activate spindle assembly factors (Carazo-Salas *et al*, 1999; Kalab *et al*, 2002) such as HURP, a bundling factor that localizes to kinetochore-attached microtubules (Silljé *et al*, 2006; Wong & Fang, 2006; Koffa *et al*, 2006). HURP accumulates more prominently in the inner region of the spindle equator (Takenouchi *et al*, 2024), potentially cooperating with KIF11 to facilitate chromosome biorientation.

In either scenario, the spindle likely exerts stronger bipolar pulling forces on chromosomes in the inner region of its equator. This idea aligns with our recent findings that the inner region of the metaphase plate has an increased risk of premature chromosome separation, a process depending on bipolar microtubule pulling forces, in aged oocytes (Takenouchi *et al*, 2024). Premature chromosome separation is a major cause of aging-associated chromosome segregation errors in oocytes (Sakakibara *et al*, 2015; Zielinska *et al*, 2015). Identifying mechanisms that regulate the spatial distribution of bipolar forces in the spindle will contribute to our understanding of the cause of egg aneuploidy.

Additionally, this study demonstrates severe meiotic defects caused by *Kif11* haploinsufficiency in oocytes. Heterozygous mutations in the *Kif11* gene have been identified in infertile patients at fertility clinics (Wu *et al*, 2024). Although several mutant forms of KIF11 have dominant-negative effects when overexpressed (Wu *et al*, 2024), our data suggest that a heterozygous loss-of-function of *Kif11* results in severe meiotic defects in oocytes and thus female infertility. Understanding the dosage-dependent role of KIF11 is therefore critical for advancing the molecular understanding of infertility and improving assisted reproductive technologies.

## Material and Methods

### Mouse

The C57BL/6 background mice were used for genetic engineering. *Spo11-*Cre mice were obtained from the Jackson Laboratory (Strain #032646 Tg(Spo11-cre)1Rsw/PecoJ) (Lyndaker *et al*, 2013). BDF1 mice were used for the experiments shown in Fig. S3. All mouse experiments were approved by the Institutional Animal Care and Use Committee at RIKEN Kobe Branch (IACUC).

### Generation of Kif11 floxed mice

The *Kif11* conditional knockout (floxed) mice (Accession No.: CDB0012E: https://large.riken.jp/distribution/mutant-list.html) were generated by CRISPR/Cas9-mediated genome editing in C57BL/6 zygotes using single-strand oligodeoxynucleotides (ssODN) as previously described (Hashimoto *et al*, 2016). The gRNA sites were designed using CRISPRdirect (Naito *et al*, 2014), and crRNA/tracrRNA and ssODN were chemically synthesized (Fasmac Co., Ltd). The floxed alleles are depicted in Fig. S1A. Genotyping PCR was performed using following primers, followed by EcoRV digestion; 5gtFW (5’-CTC CCG GTT CTC ACT GTG TC-3’) and 5gtREV (5’-TGC ACC TTA GCC ATG TAC TTT CA-3’) (WT: 731 bp, 5’-loxP: 323 and 446 bp) and 3gtFW(5’-GGC CAA GGC TGT TTC CCT AC-3’) and 3gtREV(5’-ACA GCG TTG TCA AAG CGA AA-3’) (WT: 500 bp, 3’-loxP: 322 and 218 bp).

### Mouse oocyte culture

Female mice of three to four months old were injected with 5 IU of equine chorionic gonadotropin (eCG, ASKA Pharmaceutical) or 0.1 mL of CARD HyperOva (KYUDO) to hyperovulation. Full-grown oocytes at the germinal vesicle (GV) stage were collected from the ovaries 48 hours after injection. The isolated oocytes were cultured in M2 medium containing 200 nM 3-isobutyl-1-methyl-xanthine (IBMX, Sigma) at 37°C. Meiotic resumption was induced by washing to remove IBMX. When indicated, 100 μM monastrol was used.

### Transcriptome analysis

Low input RNA sequencing was conducted using SMART-seq v4 based method following manufacturer’s protocol with slight modification in 1/4 volume reaction. Briefly, isolated full-grown oocytes at the germinal vesicle (GV) stage were washed in washing medium [0.1% PBA in PBS] and collected manually with 0.5 μL of the washing medium in 2.125 μl cell lysis buffer [10X Lysis Buffer (TAKARA), RNase Inhibitor (TAKARA), and RNase-free water (Nacalai)] in 0.2 mL 8-strip tubes. Three oocytes were collected in each tube with one or two replicates from each mouse. The collected oocytes were lysed and stored at −80°C until library preparation.

The SMART-seq v4 Ultra Low Input RNA Kit for Sequencing (Takara) was used to reverse transcribe poly(A) RNA and amplify full-length cDNA. Samples were amplified for 13 cycles in 8-strip tubes. After the cDNA was adjusted to 0.5 ng in 1.25 μL of elution buffer [10mM Tris-HCl, pH8.5], Tn5 tagmentation-based reaction was performed with 1/4 volume of the Nextera XT DNA Library Preparation Kit (Illumina) with Nextera XT Index Kit (Illumina) according to manufacturer’s protocol. Library DNA was amplified with 12 cycles of PCR and purified using 1.8× volume of SPRISelect (Beckman Coulter) and eluted into 12 μL of the elution buffer [10mM Tris-HCl, pH8.5]. Libraries were sequenced using HiseqX with 150-bp paired-end. In total, seventeen libraries were sequenced.

Hisat2 v2.2.1 was used to align the reads to the mouse genome (GRCm38) after trimming adaptor sequences and low-quality bases using fastp with the option ‘-3 -q 15 -l 15’. The resulting binary alignment/map (BAM) files were sorted using samtools. The featureCounts v2.0.3 tool implemented in the Subread software was used to generate counts of reads uniquely mapped to annotated genes using the Ensembl (release 93) annotation gtf file. In order to quantify the first exon of Kif11, as proxy for the full-length mRNA, reads were counted using the custom annotation gtf file with all Kif11 exons removed except for the exon 1. Differential expression of Kif11 between genotypes was tested with the DESeq2 package in R, which normalizes library sizes using the relative log expression (RLE) method, using the dataset of genes expressed in at least 3 samples. All possible genotype pairs were compared with Holm’s correction for multiple comparisons.

### Live imaging

Messenger RNAs were transcribed in vitro using the mMESSAGE mMACHINE T7 Kit (Invitrogen). The following mRNAs were introduced by microinjection into fully-grown GV-stage mouse oocytes: 0.4–0.6 pg H2B-mCherry, 0.5–0.7 pg mEGFP-Map4, and 0.4–0.6 pg mNeonGreen-Cep192. Live imaging was performed with a Zeiss LSM880 confocal microscope equipped with a 40x C-Apochromat 1.2NA water immersion objective lens, controlled by MyPiC (Politi *et al*, 2018). For spindle and chromosome imaging of wild-type and cKO mouse oocytes, 14 confocal z-sections (every 3 μm) of 512×512 pixel xy images were acquired every 5 minutes, whereas for control (DMSO) and inhibitor (Monastrol) treated oocyte imaging, 25 confocal z-sections (every 1.5 μm) of 256×256 pixel xy images were acquired at 5 minutes intervals. For microtubule organizing center (MTOC) imaging, 25 confocal z-sections (every 15 μm) of 512×512 pixel xy images were acquired every 12 minutes.

### 4D Image analysis

Spindle and MTOCs were reconstructed into 3D surface renderings of EGFP-MAP4 and mNeonGreen-CEP192 signals, respectively, using Imaris software (Oxford Instruments). For each time point, the generated 3D surfaces were used to calculate the volume and its position (xyz coordinate) by center of mass. For spindle shape analysis, the aspect ratio was determined by the length to width of an ellipsoid fitted to the generated 3D surface. To generate kymographs of MTOCs, chromosome positions were recorded in 3D for each time point. Principal component analysis (PCA) of chromosome positions was then performed to define the new xyz coordinates represented by the plate (PC1 and PC2 coordinates) and spindle axis (PC3 coordinates), as well as the magnitude of explained variance of each principal axis contribution (square of standard deviation; for example, complete chromosome alignment converged to explained variance of PC1, PC2, and PC3 axis as 0.5, 0.5, and 0, respectively). The 3D-reconstructed MTOCs positions were then transformed based on the new coordinates.

### Immunostaining

Oocytes were fixed with 1.6% formaldehyde in 10 mM PIPES (pH 7.0), 1 mM MgCl2, and 0.1% Triton X-100 for 30 minutes, followed by permeabilized with PBT (PBS supplemented with 0.1% Triton X-100) at 4°C overnight. The oocytes were blocked with 3% bovine serum albumin (BSA)-PBT for 2 hours and incubated at 4°C overnight with primary antibodies. The oocytes were washed with 3% BSA-PBT and then incubated with secondary antibodies and 5 µg/ml Hoechst33342 for 2 hours. The oocytes were washed again and suspended in 0.01% BSA-PBS. The oocytes were imaged under a Zeiss LSM780 confocal microscope equipped with a GaAsP detector and a 40x C-Apochromat 1.2NA water immersion objective lens. We recorded z-confocal sections (every 1 μm) of 512×512 pixel xy images to capture the entire spindle of the oocyte using LSM780.

The following primary antibodies were used: human anti-centromere antibodies (1:200, ACA, 15-234, Europa Bioproducts), a rabbit anti-Kif11 (1:500, HPA010568, Sigma). The secondary antibodies were Alexa Fluor 488 goat anti-rabbit IgG (H + L) (A11034); Alexa Fluor 555 goat anti-human IgG (H+L) (A21433). Representative single slice or Z-projection images are shown.

### Quantification of fluorescent signal intensity

Fiji (https://fiji.sc/) was used to quantify fluorescent signals. To determine the KIF11 levels at the spindle pole, the mean fluorescence intensity of KIF11 around the peak of the signal was measured. For *Kif11^fl/del^ Spo11-Cre* oocytes, KIF11 was quantified at the chromosome centers based on the microtubule enrichment in live imaging. The levels of ACA at kinetochores were also measured. The mean background intensity was obtained in the cytoplasmic region. The ratio of the value obtained from the KIF11 to that of ACA was calculated.

For the chromosome and kinetochore distribution analysis, images were reconstructed in 3D with Imaris, and the 40 kinetochore positions were manually determined. Chromosome positions were represented as the center of its two homologous kinetochores. The degree of chromosome alignment was calculated by performing the PCA on the xyz coordinates of the 20 chromosome positions as in the MTOCs analysis above. Oocytes were considered to have “aligned chromosomes” if their cumulative explained variance of PC1 and PC2 for chromosome positions was > 0.90.

## Statistical Analysis

Statistical significance was examined with R. Statistical tests, sample sizes and *p*-values are shown in figures and figure legends. When needed, Holm’s corrections were applied for correction of multiple comparisons.

## Data availability

Expression profile data generated and analyzed in this study were deposited to the NCBI Gene Expression Omnibus (GEO, www.ncbi.nlm.nih.gov/geo/) database under accession number GSE284383.

## Acknowledgements

We thank J. Ellenberg for a microscope automation macro, the imaging and animal facilities of RIKEN BDR for technical support, the Center for Advanced Technical and Educational Supports, Faculty of Agriculture, Kyushu University for part of the image analysis, and all lab members for discussions and comments. T.M. was supported by the RIKEN Special Postdoctoral Researchers Program. This work was supported by RIKEN intramural grants, RIKEN Pioneering Project “Long-Timescale Molecular Chronobiology”, JSPS KAKENHI Grant Number 23H04948 and 21H02407 to T.S.K.

## Author contributions

T.M., A.C. and T.S.K. conceived the study. T.M. performed almost all experiments and analyses. A.C. established conditional *Kif11* knockout models. H.K. performed genetic engineering. T.S.K. supervised the project. T.S.K. and T.M. wrote the manuscript with input from all authors.

## Supplementary Figures

**Figure S1:**
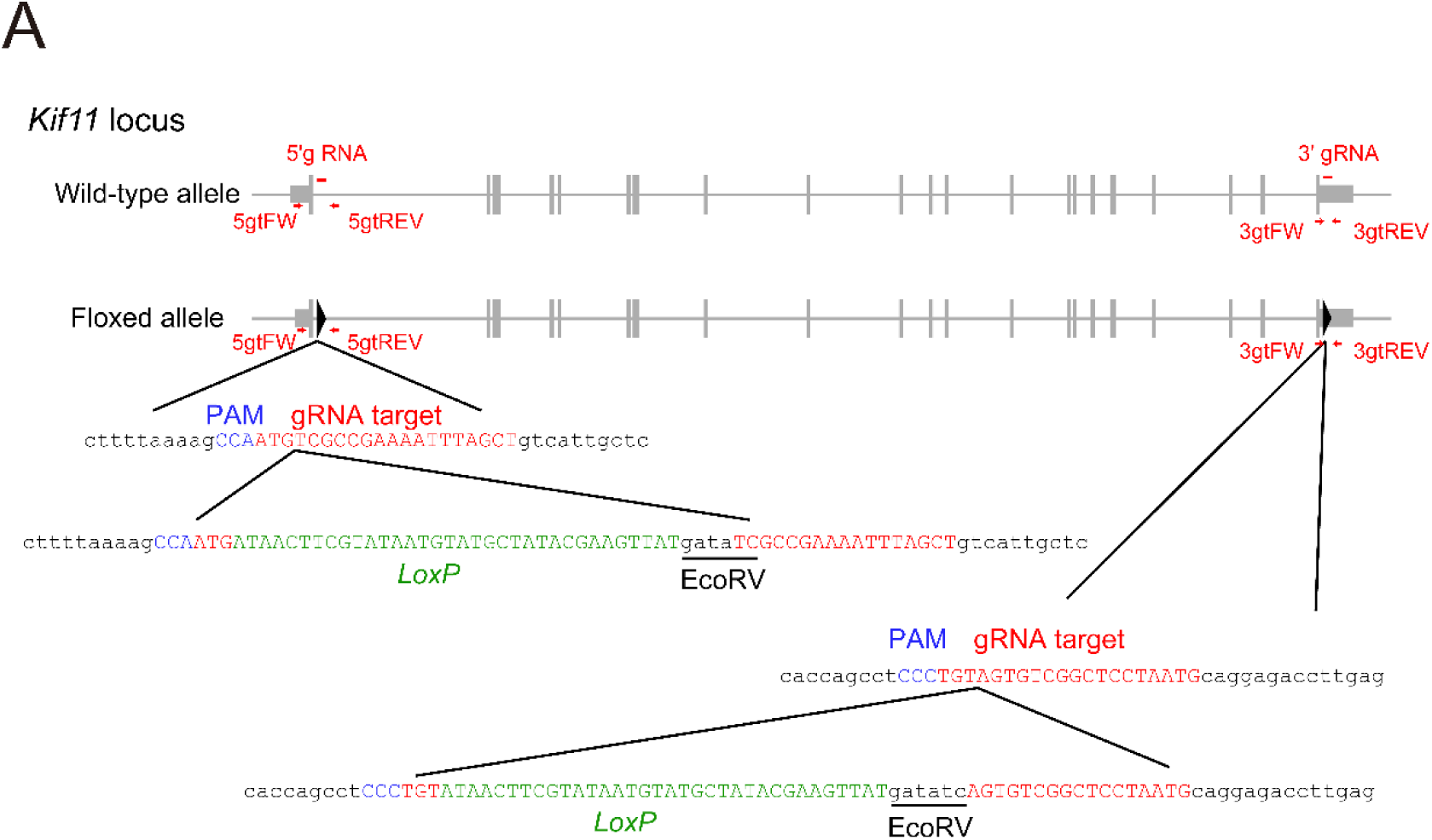
Generation of *Kif11* floxed mice. Diagram of the *Kif11* gene locus. Thin exon segments denote untranslated regions (UTR) and the thick exon segments denote coding regions. Black triangles indicate target sites for CRISPR-Cas9-mediated *Loxp* (green) insertion. Red arrows indicate primer positions used for genotyping. The protospacer adjacent motif (PAM, blue) and the guide RNA (gRNA) target sequences (red) are marked.

**Figure S2:**
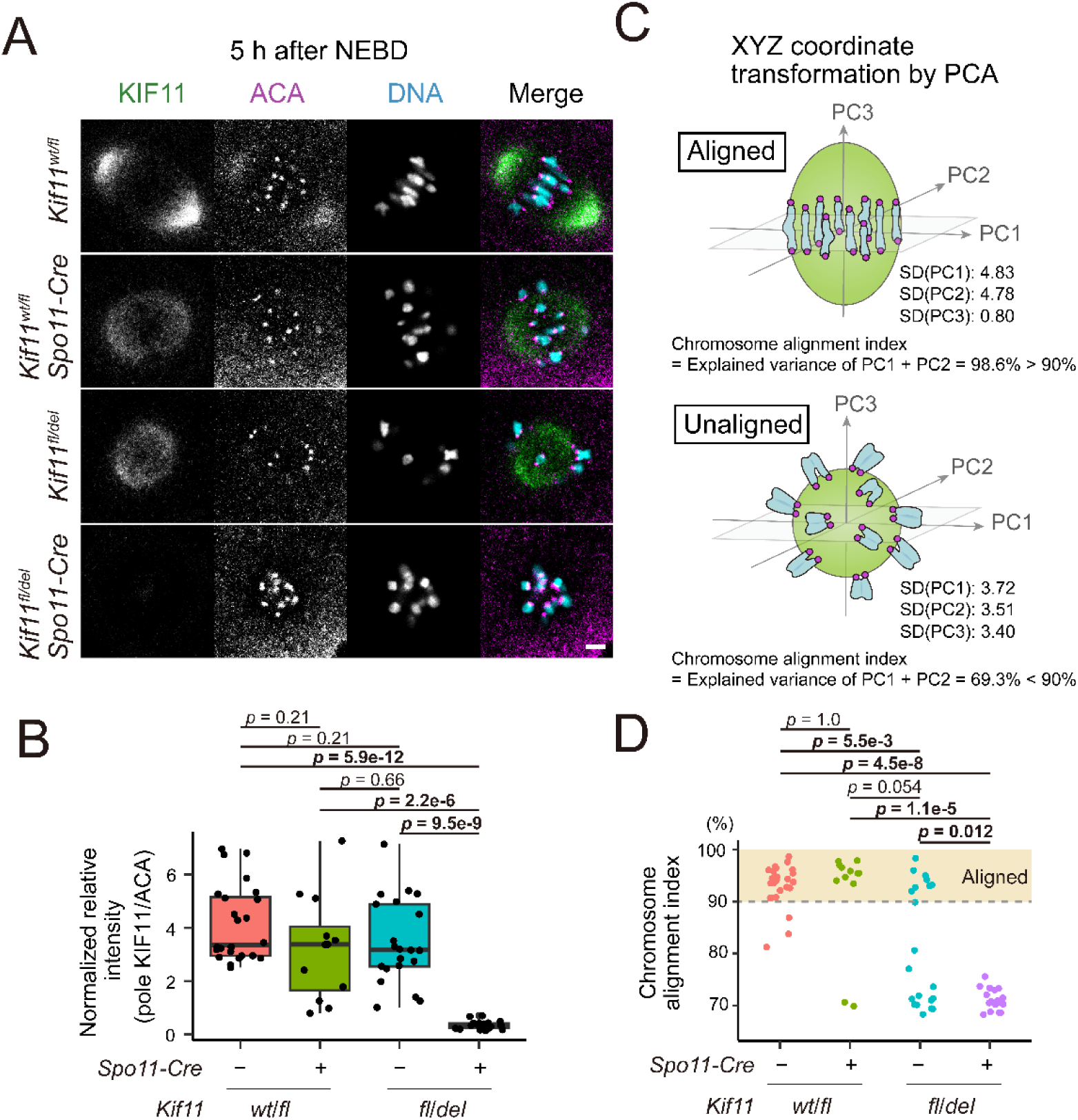
Validation of KIF11 depletion. (A) Oocytes were stained for KIF11 (green), ACA (kinetochores, magenta), and Hoechst33342 (DNA, cyan) fixed at 5 hours after NEBD (metaphase). Single z-section images are shown. Scale bar, 5 μm. (B) Quantification of KIF11 accumulated at the spindle pole. KIF11 signals in *Kif11^fl/del^ Spo11-Cre* oocytes were quantified at the chromosome centers. Welch’s t-test comparing all possible pairs with Holm’s correction for multiple comparisons. (C, D) Chromosome alignment defects in *Kif11-*deleted oocytes. Chromosome alignment index, defined as the sum of explained variance of PC1 and PC2 from PCA-transformed coordinates of chromosome position in 3D images, greater than 90% were considered as “chromosome-aligned” oocytes. The statistical significant differences (p < 0.05) in the frequency of chromosome-aligned oocytes between genotypes were compared by Fisher’s exact test comparing all possible pairs with Holm’s correction for multiple comparisons. Two independent experiments were performed.

**Figure S3:**
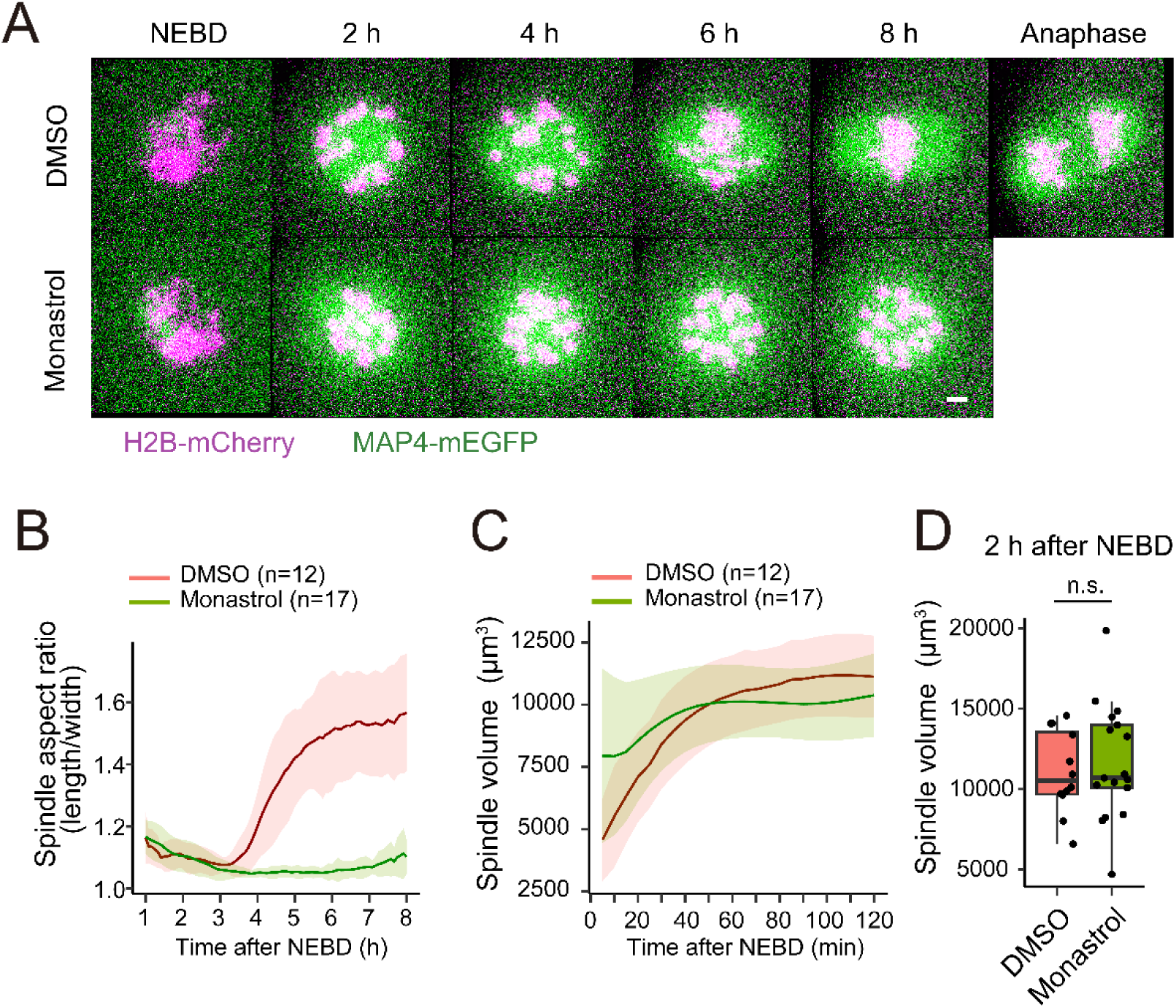
Phenotypes caused by KIF11 inhibition with monastrol. (A) Live imaging of BDF1 mouse oocytes expressing EGFP-MAP4 (microtubules, green) and H2B-mCherry (chromosomes, magenta) in the presence of DMSO (control) or monastrol. Z-projection images are shown. Scale bars, 5 µm. (B) Spindle elongation over time shown as mean ± SD. Aspect ratio (length/width) of 3D-reconstructed spindles was measured. (C) Spindle volume with mean ± SD determined from 3D-reconstructed images. (D) Box plot comparing spindle volume at prometaphase (2 hours after NEBD) between control (DMSO) and monastrol (p = 0.63, Welch’s t-test). Two independent experiments were performed.

## Supplementary Movies

**Movie EV1: KIF11 dose-dependent differences in spindle elongation.** Live imaging of spindle shape dynamics during meiosis I. Z-projection images of microtubules (EGFP-MAP4, green) and chromosomes (H2B-mCherry, magenta) are shown. Time in hh:mm. Scale bar, 5 μm. See also Fig. 2.

**Movie EV2: Difference in the spindle bipolarization in response to the dose of KIF11.** Live imaging of spindle bipolarity dynamics during meiosis I. Z-projection images of MTOCs (mNG-CEP192, green) and chromosomes (H2B-mCherry, magenta) are shown. Time in hh:mm. Scale bar, 5 μm. See also Fig. 3.

## References

Carazo-Salas RE, Guarguaglini G, Gruss OJ, Segref A, Karsenti E & Mattaj IW (1999) Generation of GTP-bound Ran by RCC1 is required for chromatin-induced mitotic spindle formation. Nature 400: 178–181

Charalambous C, Webster A & Schuh M (2023) Aneuploidy in mammalian oocytes and the impact of maternal ageing. Nat Rev Mol Cell Biol 24: 27–44

Chong MK, Rosas-Salvans M, Tran V & Dumont S (2024) Chromosome size-dependent polar ejection force impairs mammalian mitotic error correction. J Cell Biol 223: e202310010

Civelekoglu-Scholey G, He B, Shen M, Wan X, Roscioli E, Bowden B & Cimini D (2013) Dynamic bonds and polar ejection force distribution explain kinetochore oscillations in PtK1 cells. J Cell Biol 201: 577–593

Clift D & Schuh M (2015) A three-step MTOC fragmentation mechanism facilitates bipolar spindle assembly in mouse oocytes. Nat Commun 6: 7217

Drutovic D, Duan X, Li R, Kalab P & Solc P (2020) RanGTP and importin β regulate meiosis I spindle assembly and function in mouse oocytes. EMBO J 39: e101689

Dumont J, Petri S, Pellegrin F, Terret M-E, Bohnsack MT, Rassinier P, Georget V, Kalab P, Gruss OJ & Verlhac M-H (2007) A centriole- and RanGTP-independent spindle assembly pathway in meiosis I of vertebrate oocytes. J Cell Biol 176: 295–305

Hashimoto M, Yamashita Y & Takemoto T (2016) Electroporation of Cas9 protein/sgRNA into early pronuclear zygotes generates non-mosaic mutants in the mouse. Dev Biol 418: 1–9

Herbert M, Kalleas D, Cooney D, Lamb M & Lister L (2015) Meiosis and Maternal Aging: Insights from Aneuploid Oocytes and Trisomy Births. Cold Spring Harb Perspect Biol 7: a017970

Holubcová Z, Blayney M, Elder K & Schuh M (2015) Error-prone chromosome-mediated spindle assembly favors chromosome segregation defects in human oocytes. Science 348: 1143–1147

Kalab P, Weis K & Heald R (2002) Visualization of a Ran-GTP Gradient in Interphase and Mitotic Xenopus Egg Extracts. Science 295: 2452–2456

Kapitein LC, Peterman EJG, Kwok BH, Kim JH, Kapoor TM & Schmidt CF (2005) The bipolar mitotic kinesin Eg5 moves on both microtubules that it crosslinks. Nature 435: 114–118

Kapoor TM, Mayer TU, Coughlin ML & Mitchison TJ (2000) Probing Spindle Assembly Mechanisms with Monastrol, a Small Molecule Inhibitor of the Mitotic Kinesin, Eg5. J Cell Biol 150: 975–988

Kitajima TS, Ohsugi M & Ellenberg J (2011) Complete Kinetochore Tracking Reveals Error-Prone Homologous Chromosome Biorientation in Mammalian Oocytes. Cell 146: 568– 581

Koffa MD, Casanova CM, Santarella R, Köcher T, Wilm M & Mattaj IW (2006) HURP Is Part of a Ran-Dependent Complex Involved in Spindle Formation. Curr Biol 16: 743–754

Lyndaker AM, Lim PX, Mleczko JM, Diggins CE, Holloway JK, Holmes RJ, Kan R, Schlafer DH, Freire R, Cohen PE, et al (2013) Conditional Inactivation of the DNA Damage Response Gene Hus1 in Mouse Testis Reveals Separable Roles for Components of the RAD9-RAD1-HUS1 Complex in Meiotic Chromosome Maintenance. PLoS Genet 9: e1003320

Naito Y, Hino K, Bono H & Ui-Tei K (2014) CRISPRdirect: software for designing CRISPR/Cas guide RNA with reduced off-target sites. Bioinformatics 31: 1120–1123

Politi AZ, Cai Y, Walther N, Hossain MJ, Koch B, Wachsmuth M & Ellenberg J (2018) Quantitative mapping of fluorescently tagged cellular proteins using FCS-calibrated four-dimensional imaging. Nat Protoc 13: 1445–1464

Rieder CL, Davison EA, Jensen LC, Cassimeris L & Salmon ED (1986) Oscillatory movements of monooriented chromosomes and their position relative to the spindle pole result from the ejection properties of the aster and half-spindle. J cell Biol 103: 581–591

Sakakibara Y, Hashimoto S, Nakaoka Y, Kouznetsova A, Höög C & Kitajima TS (2015) Bivalent separation into univalents precedes age-related meiosis I errors in oocytes. Nat Commun 6: 7550

Schuh M & Ellenberg J (2007) Self-Organization of MTOCs Replaces Centrosome Function during Acentrosomal Spindle Assembly in Live Mouse Oocytes. Cell 130: 484–498

Silljé HHW, Nagel S, Körner R & Nigg EA (2006) HURP Is a Ran-Importin β-Regulated Protein that Stabilizes Kinetochore Microtubules in the Vicinity of Chromosomes. Curr Biol 16: 731–742

Takenouchi O, Sakakibara Y & Kitajima TS (2024) Live chromosome identifying and tracking reveals size-based spatial pathway of meiotic errors in oocytes. Science 385: eadn5529

Wong J & Fang G (2006) HURP controls spindle dynamics to promote proper interkinetochore tension and efficient kinetochore capture. J Cell Biol 173: 879–891

Wu T, Luo Y, Zhang M, Chen B, Du X, Gu H, Xie S, Pan Z, Yu R, Hai R, et al (2024) Mechanisms of minor pole–mediated spindle bipolarization in human oocytes. Science 385: eado1022

Zielinska AP, Holubcova Z, Blayney M, Elder K & Schuh M (2015) Sister kinetochore splitting and precocious disintegration of bivalents could explain the maternal age effect. eLife 4: e11389

